# Changes in DNA double-strand break repair during aging correlate with an increase in genomic mutations

**DOI:** 10.1101/2022.02.04.479125

**Authors:** Aditya Mojumdar, Nicola Mair, Nancy Adam, Jennifer A. Cobb

## Abstract

A double -strand break (DSB) is one of the most deleterious forms of DNA damage. In eukaryotic cells, two main repair pathways have evolved to repair DSBs, homologous recombination (HR) and non-homologous end-joining (NHEJ). HR is the predominant pathway of repair in the unicellular eukaryotic organism, *S. cerevisiae*. However, during replicative aging the relative use of HR and NHEJ shifts in favor of end-joining repair. By monitoring repair events in the HO-DSB system, we find that early in replicative aging there is a decrease in the association of long-range resection factors, Dna2-Sgs1 and Exo1 at the break site and a decrease in DNA resection. Subsequently, as aging progressed, the recovery of Ku70 at DSBs decreased and the break site associated with the nuclear pore complex at the nuclear periphery, which is the location where DSB repair occurs through alternative pathways that are more mutagenic. End-bridging remained intact as HR and NHEJ declined, but eventually it too became disrupted in cells at advanced replicative age. In all, our work provides insight into the molecular changes in DSB repair pathway during replicative aging. HR first declined, resulting in a transient increase in the NHEJ. However, with increased cellular divisions, Ku70 recovery at DSBs and NHEJ subsequently declined. In wild type cells of advanced replicative age, there was a high frequency of repair products with genomic deletions and microhomologies at the break junction, events not observed in young cells which repaired primarily by HR.

**Highlights:**

- Decreased DNA resection at DSBs is an early event of replicative aging
- End-joining repair increases as resection decreases at DSBs in older cells
- In older cells the products of DSB repair contain deletions and microhomologies
- DSBs associate with the NPC at the nuclear periphery more in older cells
- Old Cell Enrichment method suitable for molecular biology approaches in budding yeast

## Introduction

Age is the greatest risk factor for developing cells with genome instability and most types of cancer (Hoeijmakers 2009). There are strong links between aging and genome instability diseases. For example, DNA damage is a driver of aging and the majority of diseases with accelerated or premature aging phenotypes are caused by mutations in DNA repair factors. The overall accumulation of DNA damage is governed by the relative rate of damage formation balanced with the rate of repair, and both are impacted by age. In older cells, DNA damage formation increases and correlates with changes in genome organization and compaction (Hu et al. 2014). At the same time however, the rate of DNA repair decreases. Indeed, cells from older individuals show a lower rate of DSB repair compared to cells from younger individuals (Mayer et al. 1989; Singh et al. 1990). However, identifying the molecular underpinnings that drive age-related increases in genome instability has remained a challenge (Niedernhofer et al. 2018).

Genomic rearrangements are the types of mutations that accumulate with increased age and they arise from defects in DNA double-strand break (DSB) repair (Gorbunova & Seluanov 2016). The phosphorylation of histone H2AX (γH2AX) in humans, which corresponds to H2A (γH2A) in budding yeast, is an indirect measure of DSB formation. The level of γH2AX increases with age, yet a unified model for understanding why this occurs is proving to be complex because there are multiple pathways for repairing DSBs. There are two canonical pathways, nonhomologous end joining (NHEJ) and homologous recombination (HR), and multiple alternative (alt) pathways. The alt repair pathways are highly mutagenic compared to the canonical pathways and are used when the rates of NHEJ and HR decrease (Figure 1a). HR is considered to be error-free as it uses the sister chromatid as a repair template and can faithfully repair a DSB without loss of genetic material. By contrast, NHEJ is error-prone and often incorporates small DNA insertions or deletions at the break site. The relative use of these DSB repair pathways is not uniform across eukaryotes, or uniform in different cell types or developmental stages of the same organism, even under optimal conditions. Yeast cells repair DSBs largely by homologous recombination (HR) whereas human cells rely on NHEJ to a greater extent, however HR and NHEJ are functional in both.

**Figure 1.**
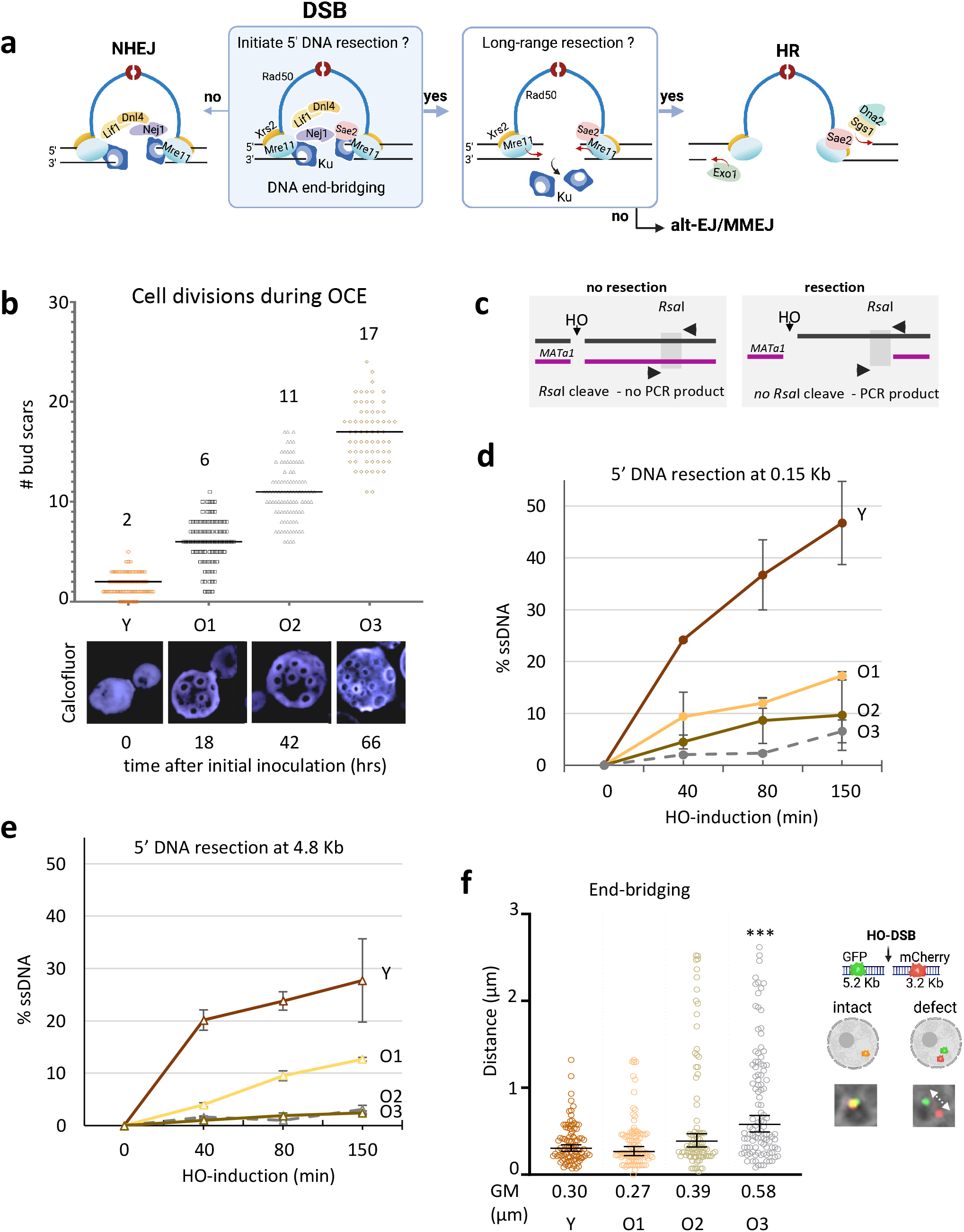
Aged yeast cells have a reduced ability to resect DSBs. **(a)** Schematic representation of DSB repair pathway choice, where repair factors play key roles in DNA end-bridging, end-resection, end-joining. If end-resection initiates then NHEJ is no longer an option. **(b)** Representative images of aging yeast cells at the different stages of old cell enrichment (OCE). Bud scars are visualized after cell staining with calcofluor. The number of bud scars is a readout for the number of cell divisions during replicative aging with images shown for young (Y) to increasing older (O1-O3) cells. Scatter plot of the average number of bud scars over 3 days of OCE in Y, O1 (18 hrs), O2 (42 hrs), and O3 (66 hours). **(c)** Schematic of the HO cut site on chromosome III showing the method used to measure the frequency of resection using the *RsaI* recognition site, the two closest sites are located 0.15 kb and 4.8 Kb away from the HO cut site. **(d and e)** Resection of DNA 0.15 Kb and 4.8 Kb away from the HO cut site was reported as the percent of single stranded DNA (ssDNA) at the indicated time points (0-150 mins) after induction of the DSB in Y and O1-O3 aged enriched cells. Analysis was performed in triplicate from at least three biological replicate experiments. **(f)** Representative image of yeast cells with tethered (co-localized GFP and mCherry) and untethered (distant GFP and mCherry) ends. Scatter plot showing the tethering of DSB ends, as measured by the distance between the GFP and mCherry foci in Y and O1-O3 aged cells 2 hours after DSB induction. The Geometric mean (GM) distance for each sample was specified under the respective sample data plot. Significance was determined using Kruskal-Wallis and Dunn’s multiple comparison test. All ages are compared to Y cells (P<0.05*; P<0.01**; P<0.001***).

Insights about how DSB repair proceeds during aging have been gained, although working with aging tissue in multicellular organisms remains extremely challenging (Gorbunova & Seluanov 2016; Niedernhofer et al. 2018). Altered intracellular distribution of Ku70/80 (Ku) and decreased protein levels for Ku and Mre11 have been reported in aging human cells (Um et al. 2003; Ju et al. 2006; Seluanov et al. 2007; Mao et al. 2012). However, changes in protein expression do not necessarily specify which pathway will be used to repair a DSB. Highlighting this case in point, the rate of HR repair decreased in the germline of old flies compared to young flies, yet the expression of HR components increased rather than decreased (Delabaere et al. 2017).

Cell cycle properties and genome maintenance factors are widely conserved in all eukaryotes and budding yeast has proven to be a robust system for identifying many of the steps in DSB repair. Computational models of aging in unicellular yeast predict that the repair of DNA damage is a critical event impacting cell fitness and replicative lifespan (Song & Acar 2019; Schnitzer et al., 2020). Moreover, recent work has shown a reduction in the levels of HR proteins in older yeast cells as well as a reduction in the use of single-strand annealing (SSA), which is an alternative repair pathway that relies on DNA resection and sequence homology (Pal et al. 2018; Young et al. 2019). However, there is relatively little known about the integrity of DSB repair mechanisms over the replicative lifespan of yeast. Much of the seminal work for monitoring physical events in DSB repair has used the galactose-inducible HO endonuclease system, which creates one site-specific DSB synchronously in the yeast genome (Connolly et al. 1988). Here, we use this well-characterized system to evaluate key steps in DSB repair in different stages of replicative aging. We measured the recruitment of HR and NHEJ repair factors to the HO-induced DSB during aging, determined that DSBs accumulate at the nuclear pore complex in older cells, and measured repair products as replicative age increased by sequencing across the break junction.

## Results

### Old cell enrichment and DSB repair pathway choice

Due to difficulties in obtaining old yeast cells, knowledge about age-related changes in DSB repair is only starting to emerge (Janssens & Veenhoff 2016; He et al. 2018; Pal et al. 2018; Young et al. 2019). We have optimized an old cell enrichment (OCE) approach with cells containing the HO-DSB where one site-specific DSB can be created by galactose-induction of the HO endonuclease (Connolly et al. 1988; Sinclair 2013). This enhanced OCE method transformed our ability to rapidly collect a large number of progressively older cells without the need for specialized microfluidics or a specific genetic background used for Mother Enrichment Program (Lindstrom & Gottschling 2009). Our method involved biotin cell labelling but differed from previous approaches by the technology used to enrich for old cells. Biotin labelled cells bind to ferromagnetic charged anti-biotin antibody coupled microbeads in a XS column (Miltenyi), which amplifies the magnetic field 10,000-fold when used in combination with the SuperMACS™ II separator system (Supplementary Figure 1a). Young cells pass through the column and biotinylated old cells are retained. After washing to remove young cell contaminants, old cells can be eluted from the XS column by withdrawing the magnetic field. We sorted young (Y) from progressively older (O) cells at the following times 18 hrs (O1), 42 hrs (O2) and 66 hrs (O3) after the initial biotin labelling. The Y cells were the daughters from the first night of old cell enrichment that did not bind to the XS column when O1 cells were recovered. O1 cells were either used for experiments or re-inoculated into fresh media and grown 24 hrs more (O2 cells). The same process was repeated with O2-aged cells to yield O3 cells (Supplementary Figure 1a).

Haploid yeast average ∼25 divisions during their lifespan (Mortimer & Johnston 1959), and the number of cellular divisions was determined by counting the number of bud scars on the surface of the mother cell visualized by calcofluor white, which labels chitin encircling the site of budding (Figure 1b; Supplementary Figure 1b) (Vrsanská et al. 1979). O3-aged cells were propagated for 66 hours and had an average of ∼ 17 bud scars (Figure 1b). It is known that the level of Sir2, a histone deacetylase, decreases with replicative age. The target of Sir2 is histone H4 K16 and older cells obtained from this OCE method showed increased H4K16 acetylation (Supplementary Figure 1c). Therefore, in addition to bud scar count, we verified that old cells displayed other known markers of replicative aging.

Two important events at a DSB that we first assessed were DNA resection and DNA end-bridging (Figure 1a). Resection is necessary for HR, but if 5’ resection initiates, then NHEJ is prevented. DNA end-bridging holds the broken ends in close proximity and provides structural support during end processing. To determine resection at the HO-DSB we used a quantitative PCR-based approach after galactose induction of the HO endonuclease. Genomic DNA was prepared at the indicated timepoints and digested with the *Rsa*I restriction enzyme followed by quantitative PCR. This method relies on a *Rsa*I cut site and there are two located at 0.15 and 4.8 Kb from the HO-DSB (Figure 1c). If resection progresses past the *Rsa*I recognition site, then the single-stranded DNA generated from resection would not be cleaved by *Rsa*I and would amplify by PCR (Ferrari et al. 2015; Mojumdar et al. 2019; Mojumdar et al. 2022b; Hohl et al. 2020). Resection in Y cells proceeds similarly to standard overnight cultures, not passed over the XS column (Supplementary Figure 1d). We were encouraged that there were no off-target effects to resection arising during the OCE process. As cellular age increased from Y to O3, resection decreased at 0.15 Kb from the break (Figure 1d), and decreased further 4.8 Kb from the break (Figure 1e). In O2 and O3 samples, this might be partly due to a decrease in DSB formation that was most pronounced 40 minutes after induction of the HO endonuclease (Supplementary Figure 1e). However, compared to the low level of resection 0.15kb from the DSB, resection was completely abrogated 4.8 Kb in O2 and O3 -aged cells (Figure 1d and e). O1-aged cells that averaged ∼ 6 divisions showed a marked decrease in resection, yet the cell has > 75% of its lifespan remaining (Mortimer & Johnston 1959). We ruled out major cell-cycle differences between Y and O1-aged cells that might account for the marked decrease in resection, which occurs in S/G2 when a sister chromatid is available. Flow cytometry before galactose induction showed that Y cells had ∼9% more cells in S/G2 compared to O1 (Supplementary Figure 1f), a minor difference that likely could not account for the 2-3-fold decrease in resection (Figure 1d and e).

End bridging was next measured in cells where both sides of the DSB were tagged with fluorescent markers. The TetO array and the LacO array were integrated 5.2 Kb upstream and 3.2 Kb downstream respectively from the DSB in cells expressing TetR^GFP^ and LacO^mCherry^ fusions, enabling us to visualize both sides of the break by fluorescence microscopy (Figure 1f). 2 hours after HO-induction the distance between the GFP and mCherry foci was similar for Y and O1-aged cells at 0.30 μm and 0.27 μm respectively and increased slightly in O2 cells (0.39 μm). In O3-aged cells, there was a significant increase in the mean distance (0.58 μm) and a wide distribution of inter-foci distance, which at the population level, was potentially a reflection of the HO cutting defects we observed in older cells (Figure 1f and Supplementary Figure 1e). Taken together, early in the aging process DNA resection decreased, however end-bridging remained intact, although it deteriorated later in replicative aging. HR is the main DSB repair pathway in yeast, however, most, if not all, studies investigating pathway choice have been performed with cultures containing mostly young cells. Thus, one possibility is that the preference for HR repair is a ‘young’ cell phenomenon that declines with age. To explore this, we were prompted to investigate the recruitment of the nucleases that mediate resection and other repair factors to the DSB during aging.

### Reduced recruitment of resection factors during aging

Mre11-Rad50-Xrs2 (MRX) and yKu70/80 (Ku) are the first complexes recruited to a DSB (Figure 2a) (Wu et al. 2008). DNA resection occurs through a two-step process (Ciccia & Symington 2016). In the first step, Sae2, the yeast homologue of human CtIP, activates Mre11 endonuclease to cleave ∼ 100 bp away from the DSB and initiate 3’-5 DNA resection towards the break. This causes Ku to dissociate, and when that happens, NHEJ is prevented. In the second step of resection, two functionally redundant 5’ to 3’ nucleases, Dna2 in complex with Sgs1 and Exo1, catalyze long-range resection (Zhu et al. 2008; Cejka 2015). The physical presence of the MRX complex is critical for all stages of resection because both long-range nucleases require MRX for their localization. (Mimitou & Symington 2008; Zhu et al. 2008). We performed chromatin immuno-precipitation (ChIP) on Mre11 and Rad50, two members of the MRX complex, with primers located 0.6 kb from the DSB (Figure 2b and c). The recovery of both increased 2 to 4-fold in O1 and O2 cells. Unfortunately, ChIP with O3 was precluded due to inefficient HO-cutting (Supplementary Figure 1e). O3 cells also showed a significant reduction in survival with continuous galactose induction (Supplementary Figure 1g). The increased recovery of Mre11 and Rad50 was surprising as previous work showed that older cells have a decreased level of HR proteins (Pal et al. 2018). Consistent with Pal et al., we also observed a decrease in the overall expression of HR proteins, including MRX components (Supplementary Figure 2a). Changes in expression levels have been used as readouts for a cell’s proficiency in a particular repair pathway (Mao et al. 2012, p.201; Titus et al. 2013; Delabaere et al. 2017). However, these results show that there is not necessarily a direct correlation between protein levels and their recruitment levels.

**Figure 2.**
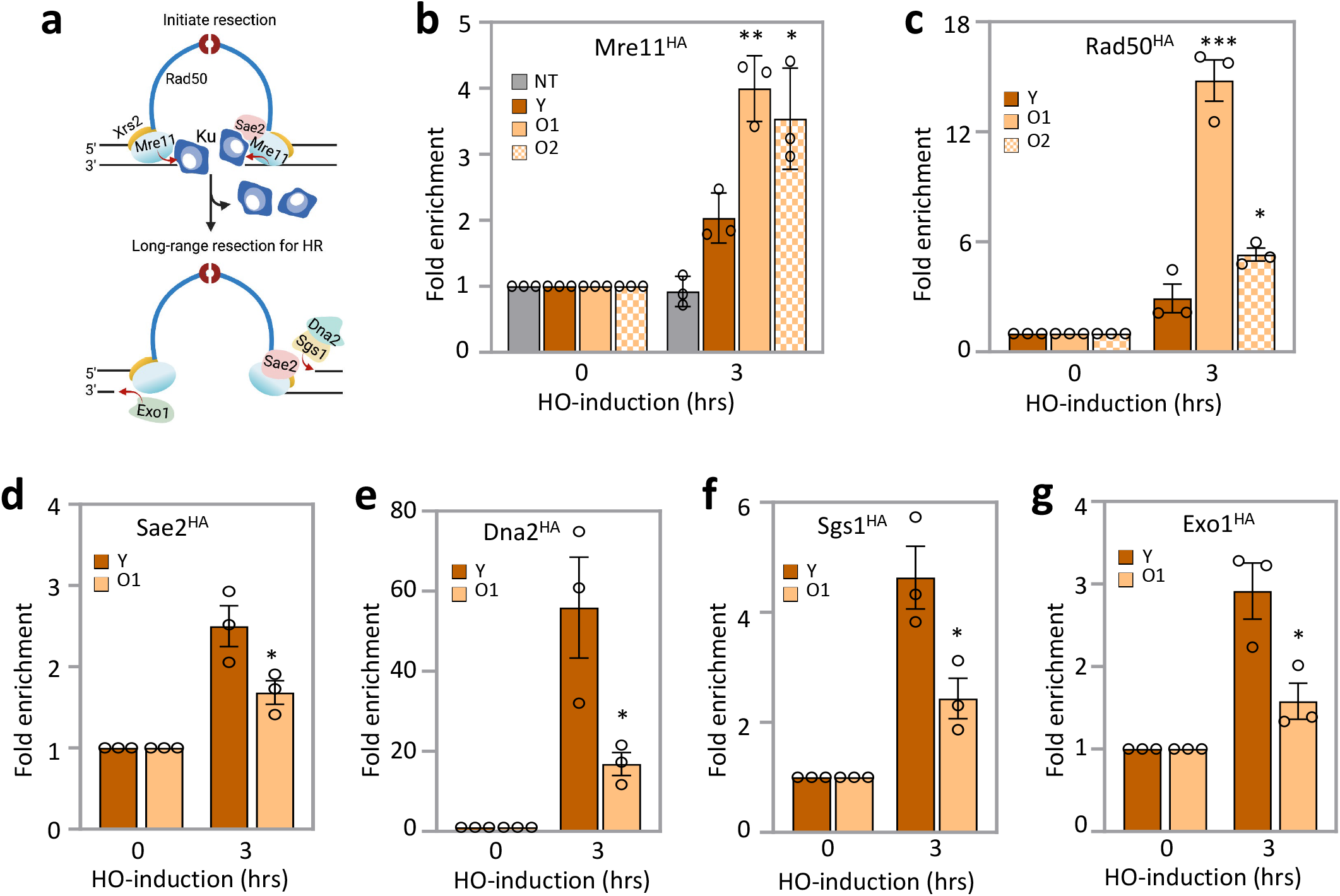
Recruitment of homologous recombination (HR) repair factors during aging. **(a)** Schematic model showing the resection steps of homologous recombination. MRX and Ku are recruited first to the DSB. Sae2 then binds Mre11 endonuclease which initiates short range resection and leads to the removal of Ku. Long-range resection is then carried out by either Exo1 or Dna2-Sgs1. **(b and c)** ChIP showed enrichment of no tag control (JC-727), Mre11^HA^ (JC-3802) or Rad50 ^HA^ (JC-3306) 0.6kb away from the DSB at 0- and 3-hour time points in Y and O1-O2 aged cells. The fold enrichment at the break was normalized to enrichment at the *SMC2* locus. **(d-g)** ChIP showed enrichment of Sae2 ^HA^ (JC-5116), Dna2^HA^ (JC-4117), Sgs1^HA^ (JC-4135), or Exo1^HA^ (JC-4869) 0.6kb away from the DSB at 0- and 3-hour time point in Y and O1 aged cells. The fold enrichment was normalized to enrichment at the *SMC2* locus. For all the experiments, the error bars represent the standard error of three replicates. Significance was determined using a 1-tailed, unpaired Student’s t test. All comparisons are to Y cells and marked (P<0.05*; P<0.01**; P<0.001***).

In contrast to MRX, the recruitment of Sae2 was reduced in O1 cells (Figure 2d), as was the recruitment of the long-range resection components, Dna2-Sgs1 and Exo1 (Figure 2e-g). In all, O1-aged cells, which have undergone ∼ 6 divisions showed a decrease in the recruitment of key factors driving DNA resection, although MRX showed increased retention. Our observations underscore the value of performing molecular and biochemical work in old cells as we gained mechanistic information about age-related changes in DSB repair. We did not perform ChIP with these factors aged to the O2 stage as we found it unlikely that their recruitment, which decreased in O1, would increase in older cells with an even greater resection defect.

### Increase in error-prone DSB repair with aging

We were curious about how DSB repair proceeded when resection, which is the main event driving HR, declined early in the aging process. Thus, we were next prompted to investigate the usage of NHEJ as end-joining reactions are supported structurally by DNA end-bridging (Hohl et al. 2015), and this remained intact from Y to O2-aged cells. As mentioned, Ku is one of the first responders to a DSB and it plays a key role in recruiting the other NHEJ factors such as Nej1 and Lif1-Dnl4 (Zahid et al. 2021). We measured the recruitment levels of the core NHEJ factors by ChIP. All the factors showed a similar level of recruitment to the DSB in Y and O1 cells, except for Ku70, which showed increased recovery in O1 cells (Figure 3a and b, Supplementary Figure 3a).

**Figure 3.**
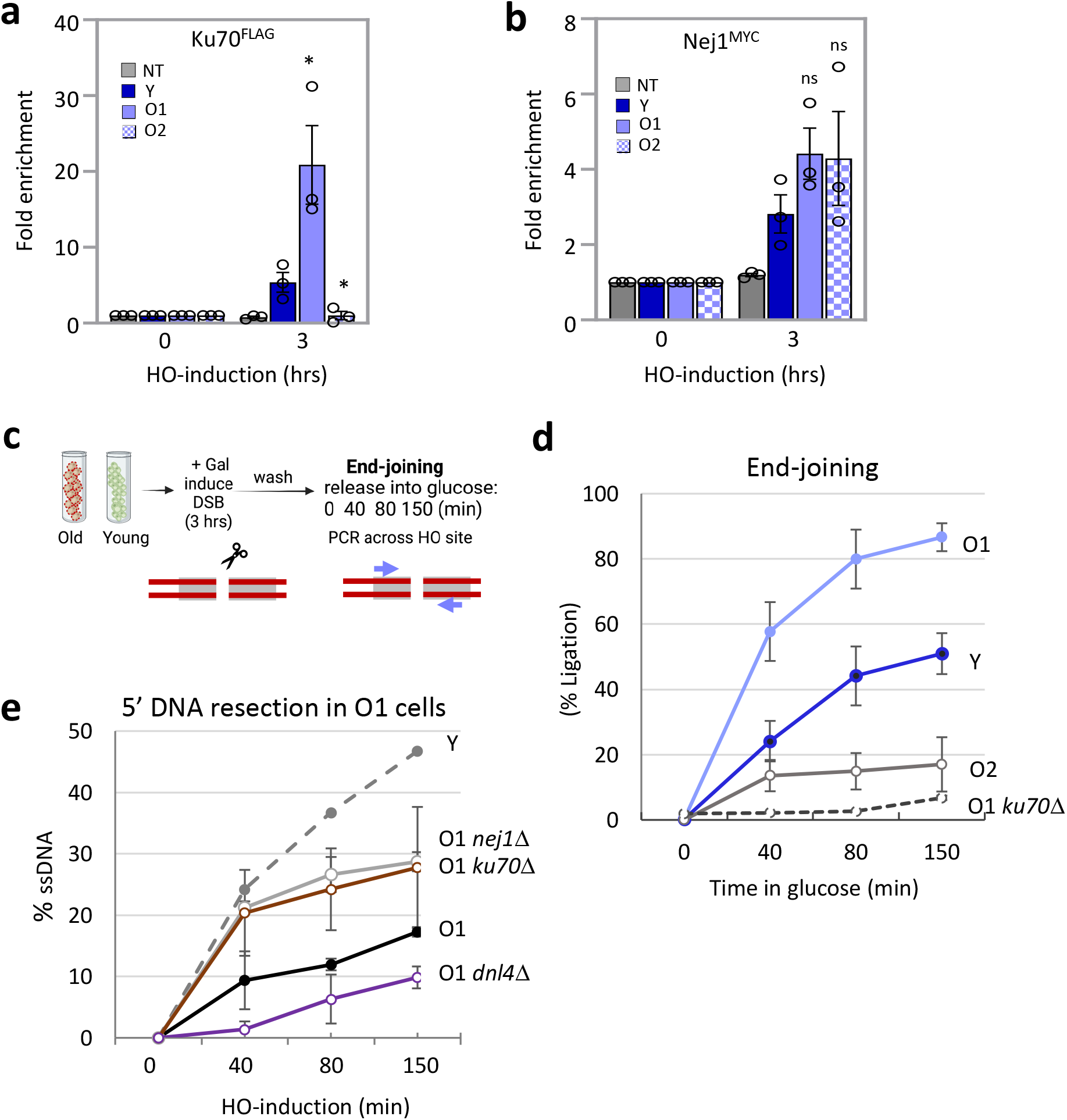
Non-homologous end joining (NHEJ) in aging cells. **(a and b)** ChIP showed enrichment of Ku70 ^Flag^ (JC-3964) and Nej1 ^Myc^ (JC-1687) 0.6kb away from the DSB at 0- and 3-hour time point in Y and O1-O2 aged cells. The fold enrichment was normalized to enrichment at the *SMC2* locus. For all the experiments, the error bars represent the standard error of three replicates. Significance was determined using a 1-tailed, unpaired Student’s t test. All comparisons are to Y cells and marked (P<0.05*; P<0.01**; P<0.001***). **(c)** Schematic of qPCR ligation assay of the HO cut site on chromosome III. Following DSB induction for 3 hours, cells were released into glucose. End-joining repair was determined with qPCR amplification with primers flanking the cut site. **(d)** Ligation was determined in WT (JC-727) and *ku70*Δ (JC-1904) in Y and O1-O2 aged. Ligation of DNA at the HO cut site is measured as the percent signal amplified 0-150 minutes following the release into glucose. **(e)** Resection of DNA 0.15kb away from the HO cut site was reported as the percent of single stranded DNA (ssDNA) at the indicated time points (0-150 mins) after induction of the DSB in Y and O1 aged cells for WT (JC-727), *nej1*Δ (JC-1342), *ku70*Δ (JC-1904) and *dnl4*Δ (JC-3290). Cut efficiency for all strains and timepoints was determined and reported in Supplementary Table 2. Analysis was performed in triplicate from at least three biological replicate experiments. For all the experiments - error bars represent the standard error of three replicates.

We next determined the level of end-ligation repair in old cells. For this, a DSB was induced by galactose for 3 hours, then cells were washed and released into glucose to prevent further re-cutting (Figure 3c). At the indicated time-points, genomic DNA was prepared followed by qPCR using primers flanking the break site. Amplification across the DSB after release into glucose would occur if the broken DNA ends were joined (Mojumdar et al. 2022a). Ligation in O1-aged cells increased 2-fold above Y cells and correlated with the increased recovery of Ku70 (Figure 3a and d). This increase in end-joining was dependent on NHEJ as deletion of *KU70* in O1 cells resulted in a complete disruption of end-joining (Figure 3d). As aging progressed to O2, there was a significant reduction in the rate of end-joining (Figure 3d), and this correlated with a marked decrease in Ku70 recovery at DSBs in O2-aged cells (Figure 3a). By contrast, the recovery of Nej1 at DSBs was not statistically different in Y-O2 aged cells (Figure 3b). It is notable that the recovery of Nej1 and MRX did not decrease as cells progress from Y to O2 as both factors function to maintain end-bridging, which remained intact at these stages of replicative aging (Figure 1f; Hohl et al. 2015; Mojumdar et al. 2019).

The relative change in 5’ resection at DSBs highlights the fluid nature of HR and end-joining repair pathway choice early in replicative aging. In O1 cells, resection decreased (Figure 1d and e) and end-joining increased suggesting that age-related declines in HR can be compensated for by an almost equivalent rise in NHEJ. We tried to increase HR by disrupting NHEJ in O1 because in Y cells resection increased when core NHEJ factors were deleted (Supplementary Figure 3c). However, unlike in Y cells, preventing NHEJ with *dnl4*Δ in O1-aged cells, did not give rise to a compensatory increase in resection (Figure 3e), suggesting changes in NHEJ per se at this stage cannot restore HR repair. Of note, deletions of *KU70* or *NEJ1* did increase resection (Figure 3e), however this is attributed to their additional roles of inhibiting Exo1 and Dna2 respectively rather than their essential NHEJ function shared with *DNL4* (Mimitou & Symington 2010; Mojumdar et al. 2019; Mojumdar et al. 2022b; Mojumdar et al. 2022a). It is striking that the increase in resection by NHEJ disruption wanes in older cells, suggesting that genotype-phenotype readouts can not only change with age, but that age can have more penetrant impact than genetic manipulations.

### During aging imprecise DSB repair increases the mutational burden on cells

Recent work in mammals, flies, and yeast has shown that difficult to repair DNA damage relocates to specific ‘repair sites’ at the nuclear periphery (Nagai et al. 2008; Chiolo et al. 2011; Chung & Zhao 2015; Ryu et al. 2015; Su et al. 2015; Churikov et al. 2016; Horigome et al. 2016). Previous work showed that DSBs relocate to the periphery within 1 hour of HO-cutting and can be detected for up to 4 hours after induction. In WT cells after galactose induction, the level of DSBs increase in the outermost zone 1 (Figure 4a) (Nagai et al. 2008; Horigome et al. 2014; Horigome et al. 2016).

**Figure 4.**
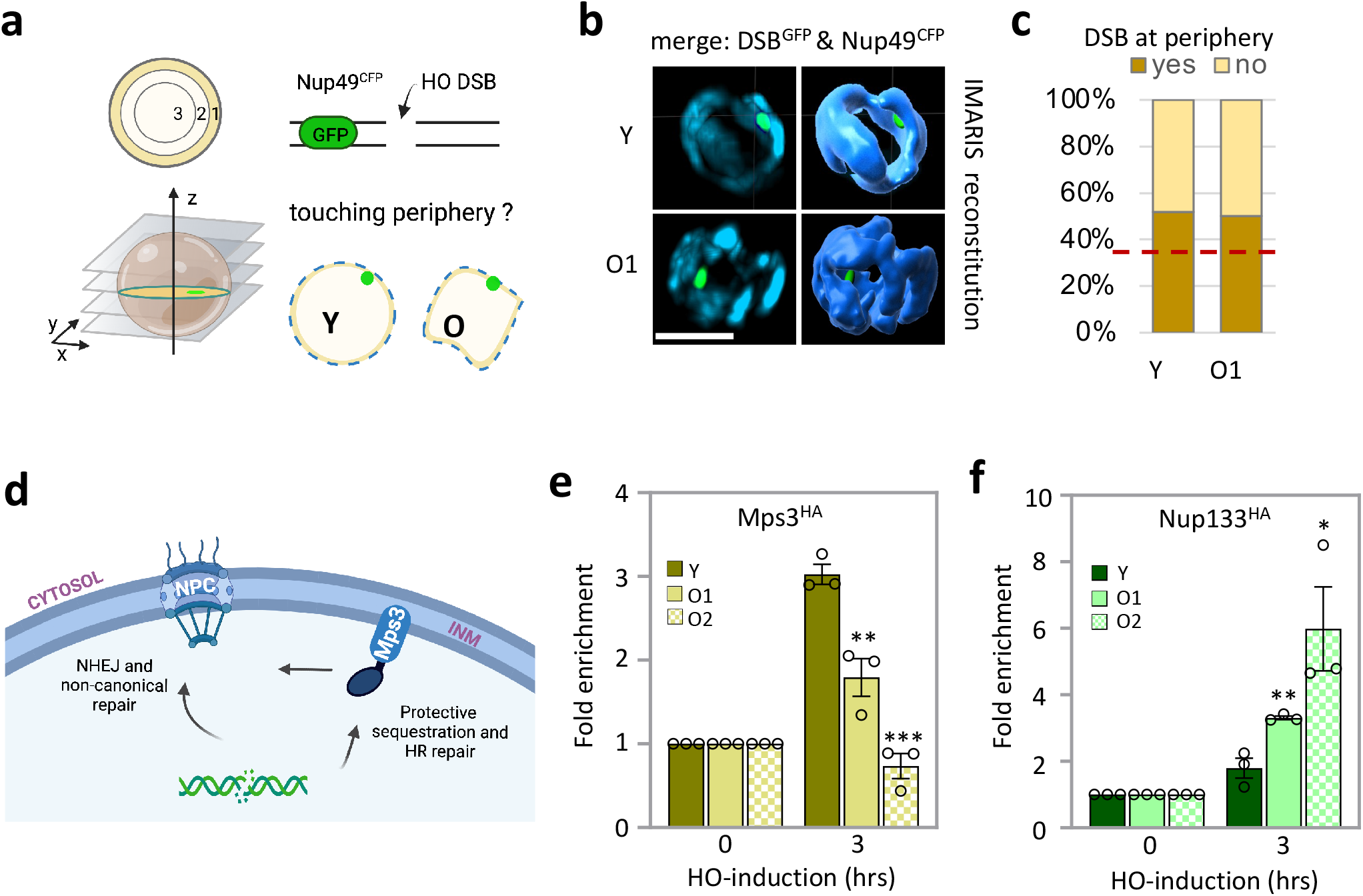
Relocation of double strand breaks (DSBs) in aged cells. **(a)** Schematic for measuring the position of the DSB at the nuclear periphery by microscopy for a GFP-tagged double strand break and CFP-tagged Nup49. Z stacks were used to determine if the DSB was in contact with the nuclear periphery. **(b)** 2D and 3D Reconstruction of the nuclear periphery using IMARIS showing the DSB ^GFP^ and Nup49 ^CFP^. **(c)** Percentage of Y and O1 aged cells where the DSB was touching the nuclear periphery. **(d)** Model for DSB re-localization to Mps3 which is located in the inner nuclear membrane (INM) of the nuclear envelope for repair via homologous recombination (HR) or to the nuclear pore complex (NPC), the site of NHEJ and non-canonical /alternative (alt)- repair pathways. **(e and f)** ChIP showed enrichment of Mps3 ^HA^ (JC-3167) or Nup133 ^Myc^ (JC-1510) 0.6kb away from the DSB at 0- and 3-hour time point in Y and O1-O2 aged cells. The fold enrichment was normalized to enrichment at the *SMC2* locus. For all the experiments, the error bars represent the standard error of three replicates. Significance was determined using a 1-tailed, unpaired Student’s t test. All comparisons are to Y cells and marked (P<0.05*; P<0.01**; P<0.001***).

Given there were age-related changes in repair pathway usage, we next investigated whether there were changes in DSB localization to the periphery. To do this, we performed high-resolution microscopy and monitored the repositioning of the HO-DSB after induction of the HO endonuclease. The DSB was visualized by expression of LacI^GFP^ in cells where an array *lacO* sites were integrated 4.4 kb from the HO site at the *MAT* locus. A focal stack of images through a field of cells allowed us to determine the position of the DSB relative to the nuclear periphery, visualized by Nup49^CFP^. The nuclear position of the DSB was determined in Y and O1 after OCE. O1 cells showed altered nuclear morphology as visualized by Nup49^CFP^ in 2D and 3D reconstructed images (cyan; Figure 4b). This prevented quantitative zoning analysis, which relies on three concentric circles representing equal areas within the nucleus. However, the overlap of GFP-CFP signals indicated that 52% and 50% of DSBs touched the nuclear periphery in Y and O1-aged cells respectively (Figure 4c), which was above the 33% level if the distribution was random. These results demonstrate that 2 hours after HO induction that irreparable DSBs in O1-aged cells were enriched at repair centers located at the nuclear envelope similarly to Y cells.

In yeast, the nuclear periphery has two independent sites where yeast damage accumulates, the nuclear pore complex (NPC) and the SUN domain protein, Mps3 (Kalocsay et al. 2009; Oza et al. 2009). These sites have different requirements for damage localization and lead to different repair outcomes. DNA intermediates with extensive resection at DSBs are targeted to Mps3, a protective environment that promotes canonical HR and reduces promiscuous recombination events in S/G2. By contrast, DSBs without extensive resection become enriched at the NPC in all cell cycle phases (Figure 4d). The nuclear pore is associated with NHEJ and non-canonical alternative repair pathways (Khadaroo et al. 2009). ChIP with NPC components and Mps3 have verified microscopy-based methods and can also quantitatively distinguish between these two specific locations at the periphery. We followed up with ChIP in O2-aged cells because we felt microscopy-based approaches posed a challenge given the age-related morphological changes, and because ChIP had worked well for measuring the recruitment of repair factors to DSBs in Y to O2 cells (Figure 2b-g, Figure 3a and b)

We performed ChIP with Nup133^Myc^, a component of the NPC, and Mps3^HA^. Consistent with previous work, DSBs in Y cells are associated robustly with Mps3 and to a lower level with the NPC after galactose induction (Figure 4e and f; Horigome et al. 2014). DSB association with Mps3 significantly declined in O1 and O2 cells, and in O2-aged cells recovery levels were indistinguishable from background (Figure 4e). By contrast, the association of DSBs with the NPC showed a progressive 2 and 3-fold increase in O1- and O2-aged cells respectively (Figure 4f). These changes were different than what was predicted when considering only age-related cell cycle alterations, and the percentage of cells in G1 and S/G2 in young and old cells (Supplementary Table 1).

As mentioned, the NPC is associated with NHEJ and other more mutagenic alt-repair pathways (Nagai et al. 2008; Khadaroo et al. 2009; Su et al. 2015; Horigome et al. 2016; Therizols et al. 2006). To determine whether age-related changes in end-joining and nuclear periphery location reflected a physiological change in how DSBs were being processed, we induced a break in young and old cells and then sequenced across the repaired break junction. Survivors of continuous HO endonuclease induction arise when the repair is mutagenic, as the recognition site is mutated, preventing re-cleavage. Consistent with previous work, the most common mutations in Y and O1 survivors were a 2-bp (+CA) insertion and a 3-bp (-ACA) deletion, and other small 1-3 bp indels (Table 1; Moore & Haber 1996; Mojumdar et al. 2019). By contrast, mutations in O2 survivors were primarily 4-6 bp deletions with 1-2 bp of microhomology (MH) (Table 1; Ma et al. 2003). Taken together, these data are consistent with a model whereby older cells use an end-joining mechanism to repair DSBs other than NHEJ, one that correlated with decreased Ku and increased NPC association, and one infrequently used in young cells.

**Table 1:**
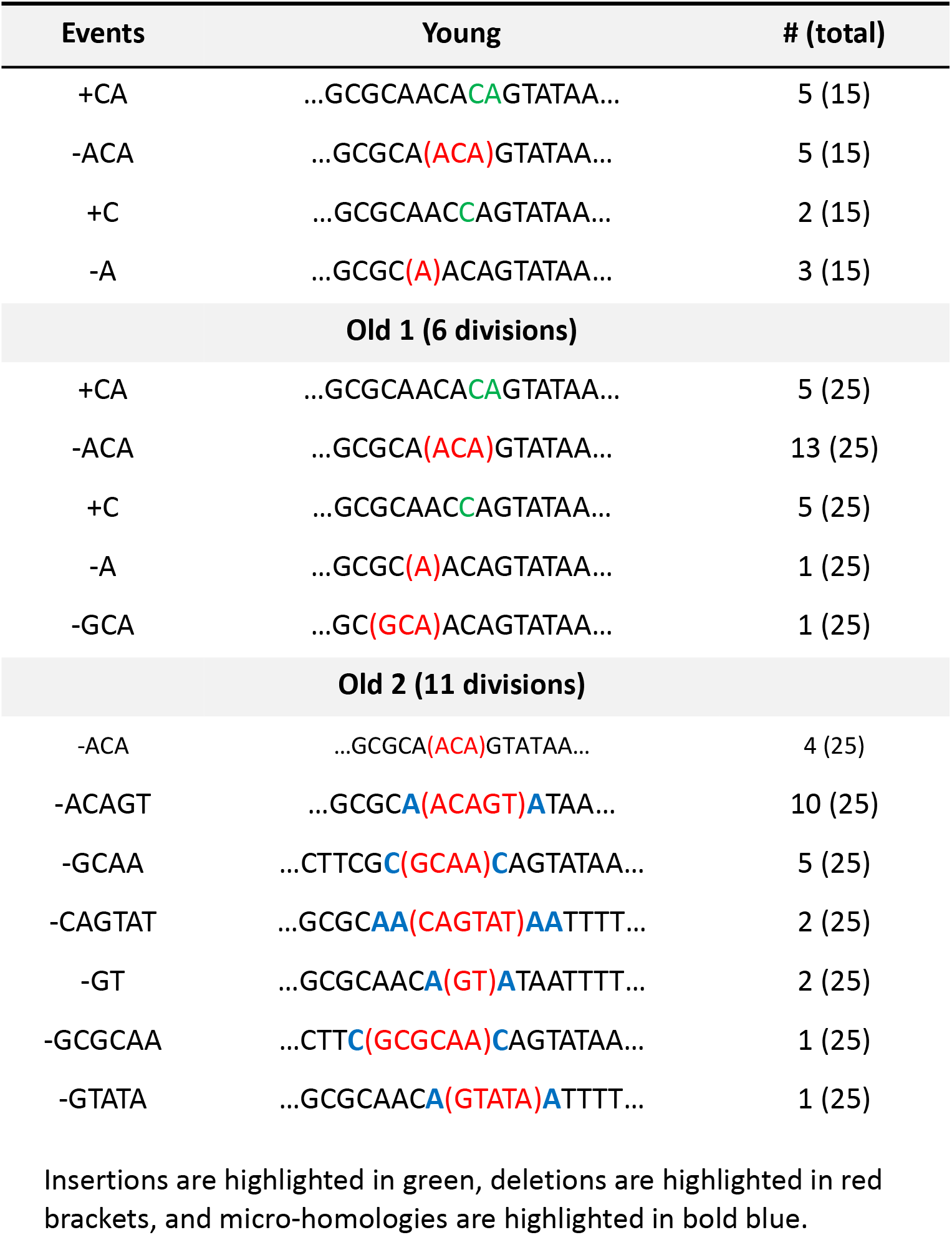
Mutation Frequency in Survivors

## Discussion

Here we report age-dependent mechanistic changes in DSB repair in the unicellular eukaryotic organism *S. cerevisiae*. Early in replicative aging, resection and the recruitment of HR factors decreased. As HR declined there was a shift towards the usage of end-joining mechanisms and a progressive increase in the frequency of repair products with larger deletions and MHs at break junctions. For this work to be possible we optimized an old cell enrichment protocol that allowed us to isolate enough old cells at various stages of aging for molecular and biochemical studies.

The old cell enrichment method used was an adapted biotin labelling method made possible by enhanced biotin binding technology that was commercially available (Supplementary Figure 1a; Sinclair 2013). Our approach was surprisingly uncomplicated and allowed large quantities of biotin labeled mother cells to be isolated with minimal young cell contamination. Many yeast aging studies have combined biotin labelling with the ‘Mother enrichment program’ (MEP), which is based on a Cre-Lox system that needs to be integrated into the yeast genome (Lindstrom & Gottschling 2009). Our approach was not combined with the MEP and did not require estradiol treatment to kill off daughter cells, but rather permitted their recovery, which might be valuable for future large-scale studies comparing daughter cells from young and old mothers.

Our work catalogued molecular changes in DSB repair in yeast cells at different stages of replicative lifespan, providing insight into how mechanisms that preserve genome maintenance in eukaryotic cells evolve with age. While the relative use and efficiency of repair pathways is known to be impacted by cell type, cell cycle phase, and ploidy, very little is known about how cellular age impacts the molecular events driving DSB repair pathway choice (Kadyk & Hartwell 1992; Trovesi et al. 2011).

DSBs are dangerous lesions and when not properly repaired they can lead to an increased mutational burden or cell death. We have measured age-dependent mechanistic changes in DSB repair and see that cellular age accentuates unfortunate repair outcomes.

In yeast, the relative use of the two canonical DSB pathways, HR and NHEJ, is 10 to 1 in favor of HR (Valencia-Burton et al. 2006). Our data suggest that this ratio is specific to very ‘young’ cells. We find that early in the cell’s replicative lifespan, by ∼6 divisions, that DNA resection at an induced DSB markedly decreased to a level that could not be attributed to slight changes in the cell cycle or in DSB cut efficiency (Figure 1d and e; Supplementary Figure 1e and f; Supplementary Tables 1-3). In cultures of the same replicative age, the frequency of end-joining increased to serve as a back-up when HR declined rather than being in competition with HR (Figure 3d and e). Deletion of *DNL4* in O1 cells did not restore resection at the HO-DSB (Figure 3e), indicating that there are intrinsic age-related decreases in HR efficiency, independent of NHEJ usage, that decline with age. These results align with previous work that showed single-strand annealing (SSA), an alternative DSB repair pathway dependent on resection, declined with increased replicative age and was not restored by *dnl4*Δ (Young et al. 2019). Decreased resection likely stems from a reduction in HR factors because their overexpression in older cells has been shown to restore DSB repair following MMS treatment and extend lifespan (Pal et al. 2018). First shown by others, and reproduced here by us, the expression of HR proteins decreased with age (Supplementary Figure 2a; Pal et al. 2018). We observed that in addition to reduced protein levels there was a significant decrease in the recruitment of specific HR factors to the DSB (Figure 2d-g). The largest relative decrease we observed between Y and O1 cells was the 3-fold decrease in Dna2 recruitment to the DSB (Figure 2e and Supplementary Figure 2a).

Our work also showed that changes in the stoichiometry of repair proteins at DSBs impacted repair pathway choice and corresponded with the shift from HR to end-joining early in replicative aging. There was a significant increase in the retention of Ku at DSBs in O1-aged cells, as well as Mre11 and Rad50 even though protein levels decreased (Figure 2c, Figure 3a, and Supplementary Figure 2a and b). Furthermore, when the recovery of most HR factors involved in resection decreased, the core NHEJ factors remained constant, including Nej1 and Lif1-Dnl4. In O2-aged cells, both resection and end-joining decreased, and this correlated with a marked decrease in Ku (Figure 3a), however the level of other end-joining factors, such as Nej1, remained constant (Figure 3b).

DNA sequencing across the break point junction showed that repair events in survivors of continuous HO induction shifted from 1-3 bp indel events when the break was induced in Y cells to larger deletions with 1-2 bp MHs at the junction when the break was induced in O2 cells (Table 1; Ma et al. 2003). It is thought-provoking to consider whether repair products in O2 cells result from NHEJ or whether these repair events can be categorized as microhomology end-joining (MMEJ). NHEJ products can have MH and one litmus test for MMEJ is whether the end-joining event occurs in the absence of Ku. MMEJ products in yeast normally have 4-6 bp MHs and occur in the absence of Ku, when the thermodynamics of NHEJ is not favored. However, most MMEJ work has been performed in young cells with reporter systems where MH length and location relative to the break have been regulated (McVey & Lee 2008; Sinha et al. 2016; Pannunzio et al. 2014).

Naturally occurring MMEJ in cells late in their replicative lifespan has never been reported. In O2 aged cells which is ∼44% of lifespan, Ku70 recovery at the DSB was reduced to background levels (Figure 3a). Thus, the local environment might indeed be a site where end-joining events are coordinated independently of Ku. Alternatively, Ku70 might be present below the level of detection. Either way, the repair products in O2 are rarely seen in young cells of the same genotype.

It is noteworthy that factors identified through the use of reporter systems to be critical for MMEJ, such as MRX and Nej1, were present at the DSB in O2 cells (McVey & Lee 2008; Sinha et al. 2016). In yeast, MRX is an integral component of most repair mechanisms, including HR and both NHEJ and MMEJ. MRX may help regulate repair pathway usage by controlling the extent of resection at the break. While HR requires extensive resection, MMEJ requires minimal resection. The 3’-5’exonuclease activity of MRX is important to initiate resection as it removes nucleotides at the DSB in ∼100 bp increments in a 3’ to 5’ direction (McVey & Lee 2008; Sinha et al. 2016). If Ku dissociates, then NHEJ cannot occur and if resection is reduced, then HR will not proceed. However, if other end-joining factors are present like Nej1 and Lif1-Dnl4 or Cdc9 ligase, then the break can be sealed through MMEJ, but with larger deletions. This represents the culmination of events in O2-aged cells.

Work performed in human and yeast have linked nuclear architecture with aging and genome stability pathways (Lans & Hoeijmakers 2006; Kubben & Misteli 2017). Naturally aged yeast showed an increase in nuclear envelope herniations associated with NPC regions and a decline in NPC assembly (Janssens & Veenhoff 2016; Rempel et al. 2020). The stability of the genome relies on the sequestration and compartmentalization of irreparable DSBs at the nuclear periphery, to environments conducive to repair. In humans, breaks at the nuclear lamina are repaired through MMEJ. In yeast, the NPC is the location where end-joining repair occurs, both NHEJ and MMEJ. In O2 aged cells there was a 3-fold increase in DBSs associated with the NPC (Figure 4f). Indeed, disruptions in nuclear pore components results in reduced survival after DNA damage exposure (Nagai et al. 2008; Loeillet et al. 2005) and single cells that maintain NPC components live longer (Rempel et al. 2020). Thus, declined NPC function in older cells might not only impact shuttling in and out of the nucleus, but it might also impact the repair efficiency of DSBs, which accumulate more at nuclear pores in older cells (Figure 4e and f).

Aging research has also focused on the rDNA repeat region (Ganley & Kobayashi 2014). The accumulation of extrachromosomal rDNA circles (ERCs), produced by recombination within the repeat region, are retained in old cells through a mechanism involving their localization to the NPC (Denoth Lippuner et al. 2014). Given this is the site where we see irreparable DSBs targeted for repair in older cells, our observations may be related to the reasons why rDNA instability and ERC formation impact global genome stability in old cells (Pal et al. 2018; Ganley & Kobayashi 2014). Increased ERCs at nuclear pores could impact the efficiency of end-joining repair in old cells, resulting in the increased rates of genomic γH2AX and translocations in old cells (Hu et al. 2014). Understanding events in the NPC compartments at the nuclear periphery might be key to understanding the relationship between aging and genome stability.

A significant number of pathways controlling yeast replicative aging appear to be fundamentally conserved, including pathways important in DSB repair and genome stability (Jasin & Haber 2016). Thus, the work presented here, which involved determining DSB repair using the HO system in yeast at different stages of replicative aging, will bring insight to the connection between cellular aging and genome maintenance in higher eukaryotes.

## Materials and Methods

All the yeast strains used in this study are listed in Supplementary Table 4 and all primers are listed in Supplementary Table 5. A resource list is listed in Supplementary Table 6. The strains were grown on various media in experiments described below.

### Old Cell Enrichment

Old cell enrichment (OCE) was based on (Sinclair 2013). For all assays 20ml of overnight culture was inoculated in 1% yeast extract; 2% peptone; 0.004% adenine sulfate dihydrate and 2% glucose (dextrose) (YPAD) medium at 25°C. Cells were then diluted to a volume of 100ml of YPAD and grown into log phase. Cells were harvested and washed three timed with 1X phosphate buffered saline (PBS) and then resuspended in 1ml PBS and mixed with 8mg of Sulfo-NHS-LC- Biotin per 1×10^8^ cells for 30 min at 30°C. Cells were washed three times with 1ml PBS and 0.1M Glycine and used to inoculate 500ml Synthetic Complete (SC) media overnight at 25°C. Cells were then harvested at 5000 RPM for 10 minutes and washed three times with PBS. Cells were then incubated in 50ml PBS and 20μl of Anti-Biotin Microbeads per 10^7^ total cells at 4°C for 30 mins. Young and old cells were then separated using a XS column on the SuperMACS II separator and washed thoroughly with PBS.

### Chromatin Immunoprecipitation

ChIP assay was performed as described previously (Mojumdar et al. 2019). Cells were collected from the OCE process and resuspended in 200ml of 1% yeast extract; 2% peptone; 2% lactate and 2% galactose (YPLG) at 30°C and cells were harvested and crosslinked at various time points using 3.7% formaldehyde solution. Following crosslinking, the cells were washed with ice-cold PBS and the pellet was stored at -80°C. Pellets were re-suspended in lysis buffer (50mM Hepes pH 7.5, 1mM EDTA, 80mM NaCl, 1% Triton, 1mM PMSF and protease inhibitor cocktail) and then lysed using Zirconia beads and a bead beater. Chromatin fractionation was performed to enhance the chromatin-bound nuclear fraction by spinning the cell lysate at 13,200 RPM for 15 minutes and discarding the supernatant. The pellet was re-suspended in lysis buffer and sonicated to yield DNA fragments (∼500 bps in length). The sonicated lysate was then incubated in beads + anti-HA or Myc conjugated dynabeads or unconjugated beads (control) for 2 hrs at 4°C. The beads were washed using wash buffer (100 mM Tris pH 8, 250 mM LiCl, 0.5% NP-40, 1mM EDTA, 1mM PMSF, protease inhibitor cocktail, and either 150 mM NaCl (for HA-epitope tagged samples) or 500mM NaCl (for Myc-epitope tagged samples). DNA that was bound by proteins was recovered by reverse crosslinking using 1% SDS in TE buffer, followed by proteinase K treatment and DNA isolation via phenol-chloroform-isoamyl alcohol extraction. Quantitative PCR was performed using the Applied Biosystem QuantStudio 6 Flex machine. PerfeCTa qPCR SuperMix, ROX was used to visualize enrichment at HO2 (0.5 kb from DSB) and HO1 (1.6 kb from DSB) and SMC2 was used as an internal control.

### Continuous DSB assay and identification of mutations in survivors

Cells from each strain were collected from the OCE process by centrifugation at 2500 RPM for 3 minutes and pellets were washed 1x in ddH_2_O and re-suspended in ddH_2_O. Cells were counted and spread on 1% yeast extract; 2% peptone; 0.004% adenine sulfate dihydrate (YPA) plates supplemented with either 2% GLU or 2% GAL. On the Glucose plates 1×10^3^ total cells were added and on the galactose plates 1×10^5^ total cells were added. The cells were incubated for 3-4 days at room temperature and colonies counted on each plate. Survival was determined by normalizing the number of surviving colonies in the GAL plates to number of colonies in the GLU plates. 100 survivors from each strain were scored for the mating type assay as previously described (Sorenson et al. 2017). Genomic DNA of sterile-type survivors was amplified with primers S1-S2, followed by DNA sequencing. All primer sequences are listed in Supplementary Table 5.

### Western Blot

Cells were lysed by re-suspending them in lysis buffer (with PMSF and protease inhibitor cocktail tablets) followed by bead beating. The protein concentration of the whole cell extract was determined using the NanoDrop. Equal amounts of whole cell extracts were added to wells of 10% polyacrylamide SDS gel. After the run, the protein was transferred to Nitrocellulose membrane at 100 V for 80 mins. The membrane was Ponceau stained (which served as a loading control), followed by blocking in 10% milk-PBST for 1hour at room temperature. The respective primary antibody solution (1:1000 dilution) was added for incubation overnight at 4°C. The membranes were then washed with PBST and left for 1 hour with secondary antibody. Followed by washing the membranes, adding the ECL substrates and imaging them.

### Resection Assay

Cells from each strain were collected from the OCE process and resuspended in 15ml YPLG. 2.5mL of the cells were pelleted as timepoint 0 sample, and 2% GAL was added to the remaining cells, to induce a DSB. Following that, respective timepoint samples were collected. Genomic DNA was purified using standard genomic preparation method by isopropanol precipitation and ethanol washing, and DNA was re-suspended in 100 mL ddH_2_O. Genomic DNA was treated with 0.005μg/μL RNase A for 45min at 37°C. 2 μL of DNA was added to tubes containing CutSmart buffer with or without the *Rsa*I restriction enzyme and incubated at 37°C for 2hrs. Quantitative PCR was performed using the Applied Biosystem QuantStudio 6 Flex machine. PowerUp SYBR Green Master Mix was used to quantify resection at *MAT*1 (0.15 kb from DSB) locus. Resection at *PRE1* was used as a negative control. *Rsa*I cut DNA was normalized to uncut DNA as previously described to quantify the % ssDNA / total DNA (Ferrari et al. 2015).

### Ligation Assay

Cells from each strain were collected from the OCE process and resuspended in 15ml YPLG. Next, 2.5mL of the cells were pelleted as ‘No break’ sample, and 2% GAL was added to the remaining cells, to induce a DSB. 2.5ml of cells were pelleted after 3hr incubation as timepoint 0 sample. Following that, GAL was washed off and the cells were released in YPAD and respective timepoint samples were collected. Genomic DNA was purified using standard genomic preparation method by isopropanol precipitation and ethanol washing, and DNA was re-suspended in 100mL ddH_2_O. Quantitative PCR was performed using the Applied Biosystem QuantStudio 6 Flex machine. PowerUp SYBR Green Master Mix was used to quantify resection at HO6 (at DSB) locus. *PRE1* was used as a negative control. Signals from the HO6 timepoints were normalized to ‘No break’ signals and % Ligation was determined.

### Bud Scar Microscopy

Following old cell enrichment 20 μl of cells were collected and fixed with 2 μl of formaldehyde. Cells were then pelleted and stored at 4°C prior to imaging. Cells are resuspended in 20 μl of PBS and stained with Calcofluor White. Cells are then placed on coverslips and imaged using Zeiss LSM 880 with Airyscan. Z-stack images were acquired with 0.35 μm along the z plane using a plan-apochromat 63x/1.40 Oil Dic M27 objective. Images were analysed using ZenBlue software. Each cell was analysed individually and >50 cells were analysed per sample.

### End-bridging Microscopy

Cells derived from the parent strain JC-4066 were collected following the OCE process. Cells were resuspended in SC with 2% lactate; 0.05% Glucose and 2% glycerol (SCLGg) media and then treated with 2% GAL for 2 hours. Cells were then washed 2 times with PBS and placed on cover slips and imaged using a fully motorized Nikon Ti Eclipse inverted epi-fluorescence microscope. Z-stack images were acquired with 200 nm increments along the z plane, using a 60 X oil immersion 1.4 N.A. objective. Images were captured with a Hamamatsu Orca flash 4.0 v2 sCMOS 16-bit camera and the system was controlled by Nikon NIS-Element Imaging Software (Version 5.00). All images were deconvolved with Huygens Essential version 18.10 (Scientific Volume Imaging, The Netherlands, http://svi.nl), using the Classic Maximum Likelihood Estimation (CMLE) algorithm, with SNR:40 and 50 iterations. To measure the distance between the GFP and mCherry foci, the ImageJ plug-in Distance Analysis (DiAna) was used (Gilles et al. 2017). Distance measurements represent the shortest distance between the brightest pixel in the mCherry channel and the GFP channel. Each cell was measured individually and > 50 cells were analyzed per condition per biological replicate.

### Nuclear Pore Localization Microscopy

Cells derived from the parent strain JC-3013 were collected following the OCE process. Cells were resuspended in SCLGg media and then treated with 2% GAL for 2 hours. Cell were collected and washed 2 times with PBS and placed on cover slips and imaged using a fully motorized Nikon Ti Eclipse inverted epi-fluorescence microscope. Z-stack images were acquired with 200 nm increments along the z plane, using a 60X oil immersion 1.4 N.A. objective. Images were captured with a Hamamatsu Orca flash 4.0 v2 sCMOS 16-bit camera and the system was controlled by Nikon NIS-Element Imaging Software (Version 5.00). All images were deconvolved with Huygens Essential version 18.10 (Scientific Volume Imaging, The Netherlands, http://svi.nl), using the Classic Maximum Likelihood Estimation (CMLE) algorithm, with SNR:40 and 50 iterations. The 2D and 3D image reconstruction was done using IMARIS microscopy image analysis software (https://imaris.oxinst.com/). When the objects assigned to CFP and GFP intensities were touching, the cell was scored as DSB located at the nuclear envelope. To measure the distance between the GFP and CFP foci, the ImageJ plug-in Distance Analysis (DiAna) was used (Gilles et al. 2017).

## Data Availability Statement

This study did not generate/analyze any code. Original data supporting the figures in the paper is available from the corresponding author on request.

## Acknowledgements

This work was supported by operating grants from CIHR MOP-82736; MOP-137062 and NSERC 418122 awarded to J.A.C. We acknowledge the resources provided by the Live Cell Imaging Laboratory. The Nikon Ti Eclipse inverted epi-fluorescence microscope system was purchased with funds from the International Microbiome Centre, which is supported by the Cumming School of Medicine at University of Calgary, Western Economic Diversification (WED) and Alberta Economic Development and Trade (AEDT), Canada.

## Authors Contributions

**Aditya Mojumdar:** Methodology, Investigation, Supervision, Formal analysis, Writing-original draft and revisions, **Nicola Mair:** Methodology, Investigation, Writing-original draft and revisions, **Nancy Adam:** Investigation, **Jennifer A Cobb**: Supervision, Conceptualization, Funding acquisition, Writing-original draft and revisions,

## Figure legends

**Supplementary Figure S1.**
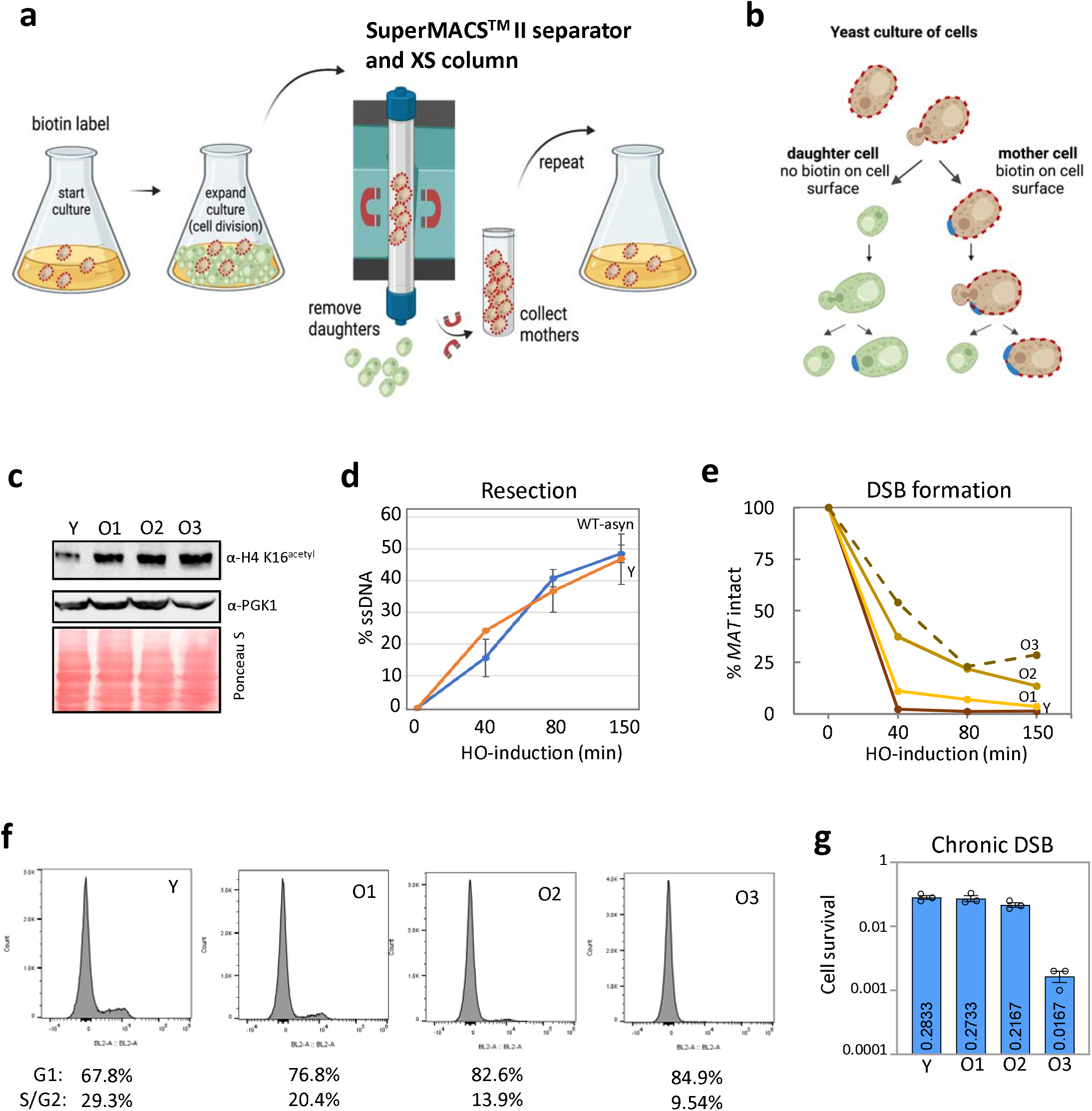
The Process of Old Cell Enrichment. Related to Figure 1. **(a)** Schematic of old cell enrichment (OCE) used to collect Y and O1-O3 aged cells. Cells were biotin labelled and used to inoculate an overnight culture of SC media. Young daughter and biotin labelled mother cells in the culture are separated over a SuperMACS™ magnet. The mother cells were then reinoculated in fresh SC media to promote further replicative aging. **(b)** Schematic depicting the biotin labelling (red dashed outline) of mother cells. The biotin labelling remains within the cell wall of only the mother because the cell wall of daughter cells have new cell wall material. **(c)** Western blot of WT cells (JC-727) showing the expression of H4K16 acetylation in Y and O1-O3 cells. PGK1 was used as a loading control. **(d)** Resection of DNA 0.15kb away from the HO cut site is measured as the percent of single stranded DNA (ssDNA) at 0, 40, 80 and 150 minutes following the induction of the DSB in WT (JC-727) overnight cultures or Y cells obtained from the OCE process. **(e)** WT cells (JC-727) aged using the OCE process to sort Y and O1-O3. The efficiency of the HO cut was determined by measuring the percentage of MAT that remains uncut (intact) at 0, 40, 80 and 150 minutes after induction with galactose. **(f)** Flow cytometry was perform with the fraction of cells in G1 and S/G2 cell cycle phase shown for Y and O1-O3 aged cells. **(g)** Percentage cell survival upon chronic HO induction in Y and O1, O2, O3 -aged cells.

**Supplementary Figure S2:**
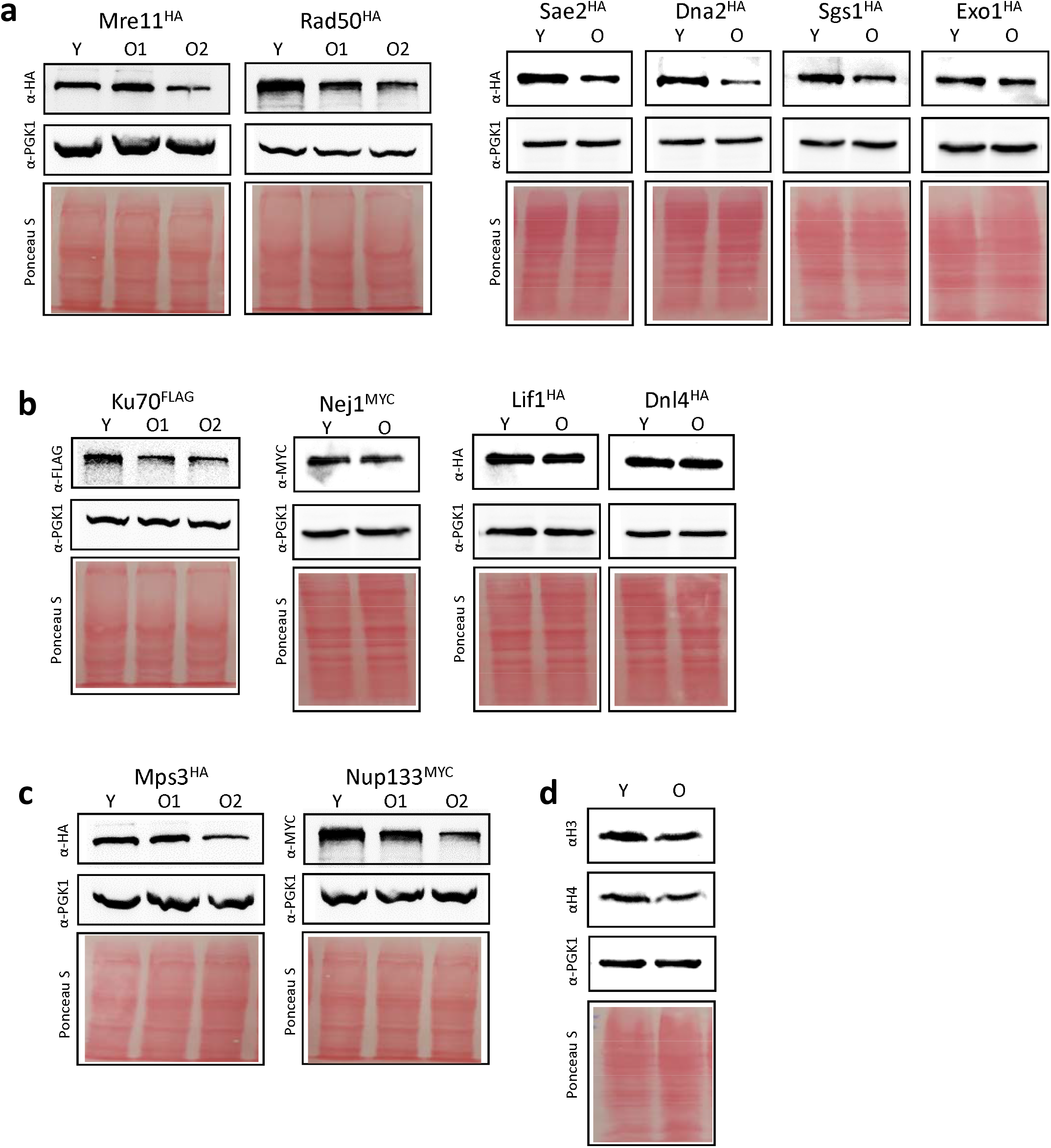
Western blot analysis of DSB repair factors. Related to Figure 2, 3 and 4. **(a-c)** Expression of DSB repair proteins in young and old cells determined by western blots with either αMyc (ab32) or αHA (F-7). The strains used were Mre11^HA^ (JC-3802), Rad50^HA^ (JC-3306), Sae2^HA^ (JC-5116), Dna2^HA^ (JC-4117), Sgs1^HA^ (JC-4135), Exo1^HA^ (JC-4869), Ku70^FLAG^ (JC-3964), Nej1^MYC^ (JC-1687), Lif1^HA^ (JC-3319), Dnl4^HA^ (JC-5672), Mps3^HA^ (JC-3167), Nup133^MYC^ (JC-1510), and (d) WT (JC-727) was used for histone H3 and H4. PGK1 and Ponceau staining were used as a loading controls.

**Supplementary Figure S3:**
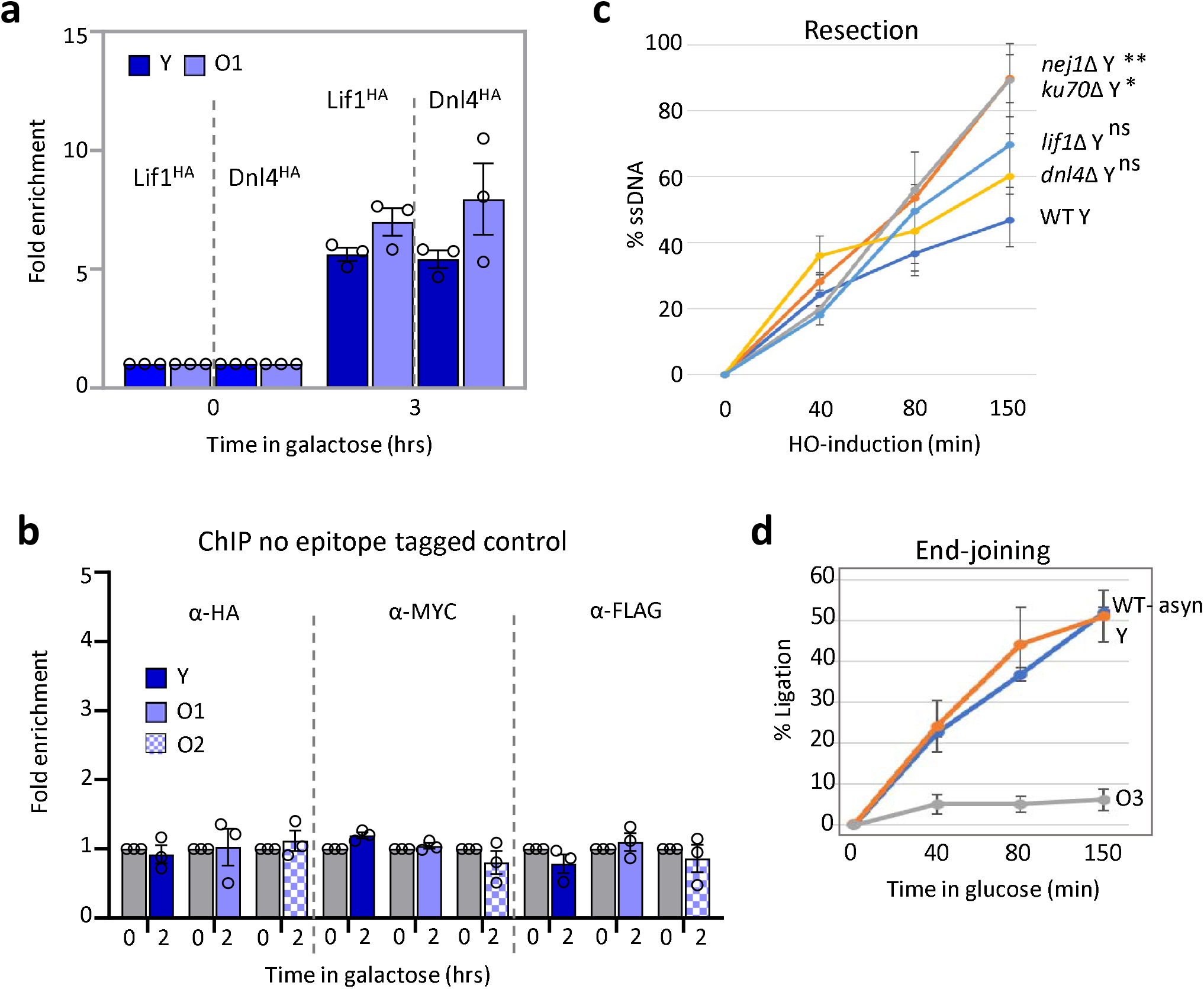
Supporting information for Figure 3. **(a)** Enrichment of Lif1^HA^ and Dnl4^HA^ (JC-3319 and JC-5672) 0.6kb away from the DSB at 0- and 3-hour time point in Y and O1-aged cells. The fold enrichment is normalized to the *SMC2* locus. **(b)** Enrichment of No tag control (JC-727) with αHA, αMYC and αFLAG antibody, 0.6kb away from the DSB at 0- and 3-hour time point in Y and O1-O2 aged cells. The fold enrichment is normalized to the *SMC2* locus. **(c)** Resection of DNA 0.15kb away from the HO cut site was reported as the percent of single stranded DNA (ssDNA) at the indicated time points (0-150 mins) after induction of the DSB in Y cells for WT (JC-727), *nej1*Δ (JC-1342), *ku70*Δ (JC-1904) and *dnl4*Δ (JC-3290). Analysis was performed in triplicate from at least three biological replicate experiments. For all the experiments - error bars represent the standard error of three replicates. Significance was determined using 1-tailed, unpaired Student’s t test. All strains compared to Y cells and marked (P<0.05*; P<0.01**; P<0.001***). **(d)** % Ligation at the HO cut site is measured as the percent signal amplified 0-150 minutes following the release into glucose. Wild type (WT) cells (JC-727) without any processing, and Y and O3 –aged cells that went through the OCE process.

